# PCDH15 Dual-AAV Gene Therapy for Deafness and Blindness in Usher Syndrome Type 1F

**DOI:** 10.1101/2023.11.09.566447

**Authors:** Maryna V. Ivanchenko, Daniel M. Hathaway, Eric M. Mulhall, Kevin T. Booth, Mantian Wang, Cole W. Peters, Alex J. Klein, Xinlan Chen, Yaqiao Li, Bence György, David P. Corey

## Abstract

Usher syndrome type 1F (USH1F), resulting from mutations in the protocadherin-15 (PCDH15) gene, is characterized by congenital lack of hearing and balance, and progressive blindness in the form of retinitis pigmentosa. In this study, we explore a novel approach for USH1F gene therapy, exceeding the single AAV packaging limit by employing a dual adeno-associated virus (AAV) strategy to deliver the full-length PCDH15 coding sequence. We demonstrate the efficacy of this strategy in mouse USH1F models, effectively restoring hearing and balance in these mice. Importantly, our approach also proves successful in expressing PCDH15 in clinically relevant retinal models, including human retinal organoids and non-human primate retina, showing efficient targeting of photoreceptors and proper protein expression in the calyceal processes. This research represents a major step toward advancing gene therapy for USH1F and the multiple challenges of hearing, balance, and vision impairment.

## Introduction

Usher Syndrome Type 1F (USH1F) is a debilitating genetic condition in which patients are born without the senses of hearing or balance and further experience gradual deterioration of vision^1-7^. The disparate symptoms of USH1F result from dysfunction in a single gene which encodes the extracellular adhesion protein protocadherin-15 (PCDH15)^5,8^. In the cochlea, PCDH15 is essential for mediating hair-cell mechanotransduction, accounting for the dramatic loss of hearing and balance associated with its mutation ^9–11^. Vision impairment in USH1F is due to retinitis pigmentosa, a progressive peripheral-to-central degeneration of the rod and cone photoreceptors of the retina ^12–14^. Most USH1F patients experience loss of night vision in their teens, which eventually progresses to tunnel vision and complete blindness^15^. The loss of vision is particularly devastating because patients rely on vision to compensate for the impairment of hearing and balance, resulting in unmet needs for gene therapy. Furthermore, the delayed manifestation of these symptoms presents a therapeutic window during which intervention may be effective, prior to photoreceptor death.

Over the last 20 years, we and others have explored the role of PCDH15 in the cochlea, elucidating its importance for mechanotransduction and hair cell development ^5,6,9,10,16–22^. However, the role of PCDH15 in the retina has remained less clear. While mouse models of USH1F faithfully recapitulate auditory and vestibular defects, they exhibit only subtle changes in vision function and no obvious change in retinal morphology^12^. Studies in amphibians and nonhuman primates (NHPs) have indicated that PCDH15 is a component of the calyceal processes^23,24^ which surround the bases of photoreceptor outer segments and maintain their integrity. Importantly, calyceal processes are absent from mouse photoreceptors, which may explain the limited visual phenotype of mouse models of USH1F ^23^.

Clinically, USH1F is an attractive candidate for intervention by gene therapy: subretinal AAV therapy was approved for clinical use^25^ and many other retinal gene therapies are in development^26^. However, the large coding sequence of PCDH15 constrains an AAV-based gene delivery. Recently, we effectively circumvented this constraint in the cochlea using two gene therapy approaches: a mini-gene approach^27^ and a base-editing approach^28^. For those studies, we generated two novel mouse models of USH1F cochlear pathology, a *Pcdh15* floxed mouse with recombination induced by *Myo15-Cre*, and a humanized mouse line bearing the common pathogenic R245X point mutation flanked by the human coding sequence. In this study, we use these models as well as human retina organoids and a primate retina to test a third circumvention of the length constraint: dual-AAV gene delivery.

Dual-AAV gene delivery relies on co-transduction of a single cell with two different viruses, each carrying half of the genetic cargo, with subsequent intracellular recombination. Here, we employ a hybrid dual AAV strategy that incorporates both trans-splicing and overlapping elements to maximize the proper concatemerization of the viral DNA^29–33^. In recent years, this strategy has been used to deliver larger transgenes to the cochlea, such as OTOF ^31,34^ and MYO7A^35^, and to the retina, such as ABCA4^36^ and MYO7A^37^. Using our mouse models, we demonstrate remarkable efficacy in correcting hearing and balance defects. For potential retinal therapy, we tested the dual-AAV therapy in two clinically relevant models: human retinal organoids and subretinal injection in the green monkey, an old-world primate. In both primate species, we demonstrate safe and efficient targeting of photoreceptors, as well as proper protein expression and localization to the calyceal processes. Our study offers promise for correcting USH1F-associated defects in hearing, balance and vision.

## Results

### Generation of dual AAV vectors and verification in HEK 293 cells

For testing gene replacement therapy in the USH1F mouse inner ear, we used the mouse *Pcdh15-CD2* isoform (NM_001142742.1), previously confirmed to be necessary for PCDH15 function in hair cells in mice ^17,38^. *Pcdh15-CD2* has a ∼5.3 kb coding sequence, too large to fit in a single AAV capsid. We engineered dual-AAV hybrid vectors so that each vector encoded about half of the full-length protein. Vector 1 (AAV-Pcdh15 5’) included a CMV promoter, the N-terminal half of the *Pcdh15* coding sequence, and a splice donor (SD) site followed by a highly recombinogenic sequence from F1 phage (AK)^32^. Vector 2 (AAV-Pcdh15 3’) included the AK sequence, a splice acceptor (SA), the C-terminal half of the *Pcdh15* coding sequence, and other regulatory elements (**Fig. 1a**). Once AAV vectors are in the same cell, the reassembly is mediated by homologous recombination of the AK sequence and/or non-homologous end joining of the inverted terminal repeats (ITRs). SD and SA sites facilitate the excision of the ITRs via trans-splicing, producing a full-length *Pcdh15* mRNA and protein (**Fig. 1a**)^30^ ^39^. This method has worked well for the dual-AAV expression of otoferlin in the cochlea ^31,34,40^. Each recombinant vector was packaged in the AAV9-PHP.B capsid, which efficiently transduces both inner hair cells (IHCs) and outer hair cells (OHCs) of the cochlea ^41–43^. We also made a second N-terminal vector that includes a hemagglutinin (HA) epitope tag at the N-terminus of PCDH15 (AAV-HA.Pcdh15 5’). The N-terminus is a short helix extending away from the first EC domain and away from the bond with cadherin-23 (CDH23), so the HA tag itself does not interfere with normal binding ^27^.

**Fig. 1:**
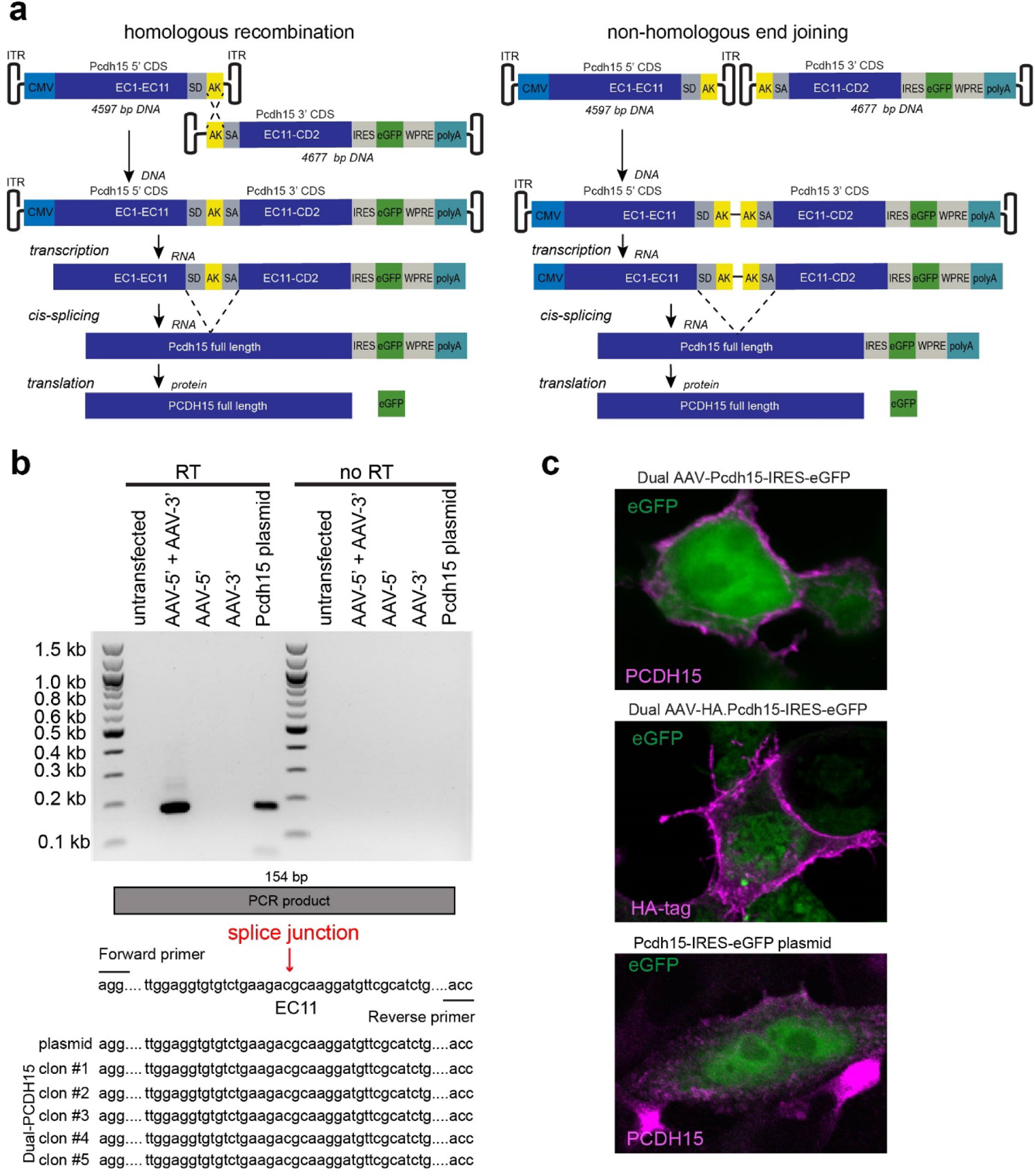
Splicing and protein production *in vitro*. **a** Recombination and splicing of dual vectors to produce full-length protein. The reassembly is mediated by non-homologous end joining of the inverted terminal repeats (ITRs) and/or homologous recombination of the highly recombinogenic (AK) sequence. The AK sequence promotes homologous recombination between the two vector DNAs. After transcription, splicing can occur from the splice donor site (SD) in vector 1 to the splice acceptor site (SA) in vector 2, creating a full-length PCDH15 mRNA. **b** mRNA was obtained from HEK cells and reverse transcribed. cDNA library was Sanger-sequenced around the splice junction, confirming proper recombination and splicing. **c** Dual-AAVs (with or without an N-terminal HA tag) were added to HEK 293 cell cultures. Immunostaining was performed with anti-PCDH15 or anti-HA antibodies (magenta). Representative confocal images demonstrate normal trafficking of PCDH15, either with or without the N-terminal HA tag, to the cell membrane after dual-AAV delivery. Labeling was appeared similar as with transfection of a single plasmid encoding full-length PCDH15. CMV, cytomegalovirus; WPRE, woodchuck hepatitis virus post-transcriptional regulatory element; polyA, polyadenylation signal; IRES, internal ribosome entry sites.

We first assessed splicing and protein production *in vitro*. An HEK 293 cell line was transduced with either AAV-Pcdh15 5’ + AAV-Pcdh15 3’ or AAV-HA.Pcdh15 5’ + AAV-Pcdh15 3’. Cells were collected for mRNA or fixed for immunofluorescence. mRNA harvested from HEK293 cells was reverse-transcribed, and a cDNA library was Sanger-sequenced around the splice junction, confirming proper recombination and splicing (**Fig. 1b**). We labeled fixed cells with antibodies to PCDH15 or to the HA tag. If, and only if, cells were transduced with both AAVs, we observed strong labeling of the plasma membranes with either anti-PCDH15 or anti-HA (**Fig. 1c**). Labeling was the same as with transfection of a single plasmid encoding full-length PCDH15, indicating that recombination and splicing were efficient.

### Dual-AAV mediated Pcdh15 coding sequence delivery restores auditory and vestibular function in an USH1F mouse model and shows no toxicity for hearing

To test efficiency of dual-AAV delivery in rescue of hearing, we used *Pcdh15^fl/fl^* conditional knockout mice, in which exon 31—encoding PCDH15’s transmembrane domain and part of the MAD12 domain—was flanked by loxP sites^27^. Exon 31 deletion by recombination occurred in hair cells when mice also carried a Cre recombinase driven by the late-onset, hair-cell-specific *Myo15* promoter ^44^.

As previously shown *Pcdh15^fl/fl^, Myo15-Cre ^+/-^* mice at 5 weeks age lacked any identifiable auditory brainstem response (ABR) to loud sound stimulation across the full frequency spectrum tested, indicating profound hearing loss^27^. Distortion-product otoacoustic emission (DPOAE) responses at 5 weeks age, diagnostic of OHC function, were largely absent at all sound intensities and frequencies tested. Scanning electron microscopy and phalloidin staining of actin revealed severely disrupted bundles: middle and short row stereocilia of IHCs and OHCs in the *Pcdh15^fl/fl^, Myo15-Cre ^+/-^* mice were shortened or totally missing. Hair cells also showed no detectable FM1-43 loading in adults, confirming no functional mechanotransduction.

AAV vectors were injected into cochleas of *Pcdh15^fl/fl^, Myo15-Cre ^+/-^* conditional knockout mice at P1 via the round window membrane (RWM), with a dose 5.0×10^10^ VGC of each vector (1.0×10^11^ VGC total). At 5 weeks, the treated animals were tested physiologically to determine the extent of hearing rescue, and cochlear tissue was processed for histological examination (**Fig. 2a**).

**Fig. 2.**
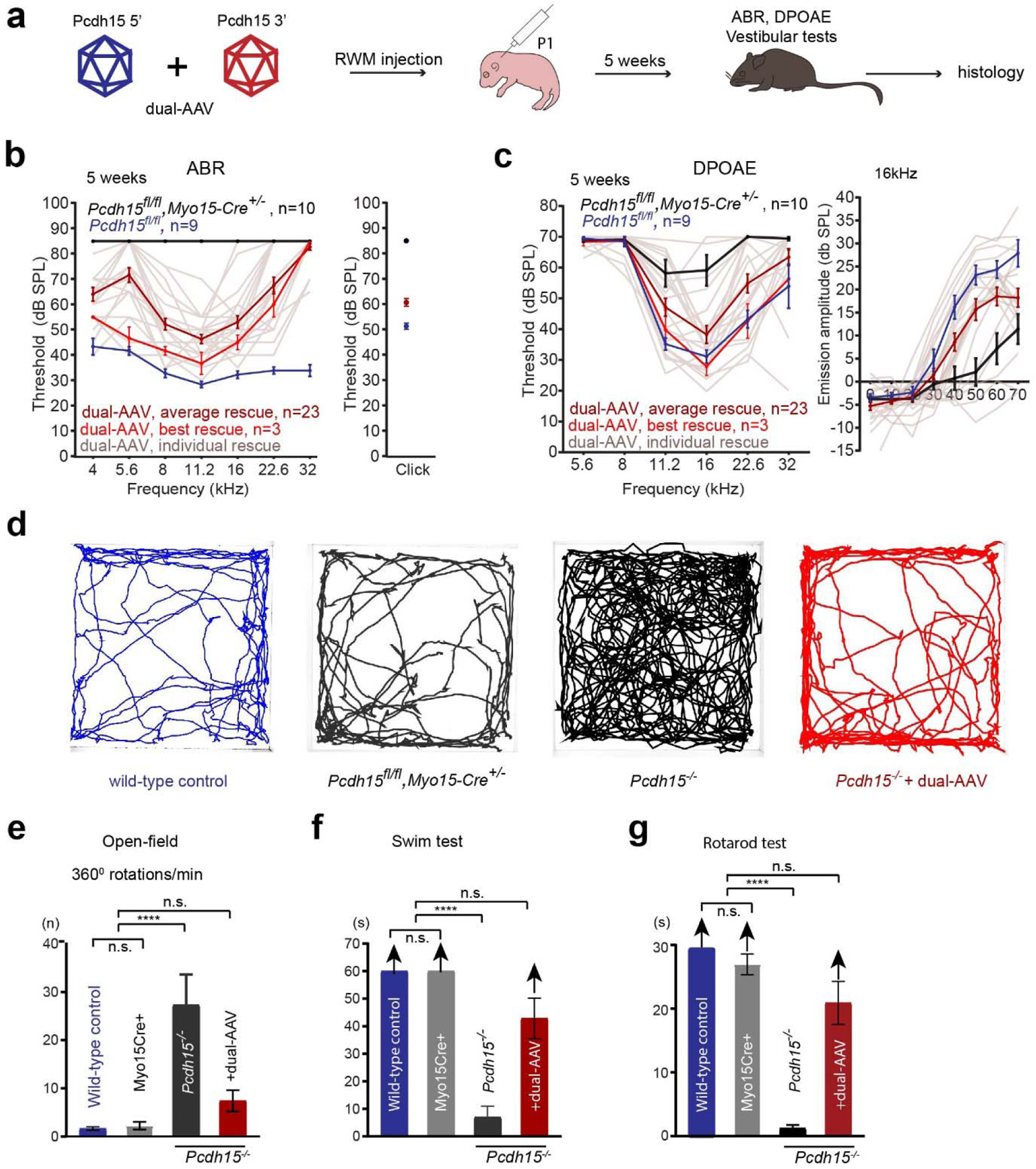
Dual-AAV delivery of PCDH15 successfully restored hearing in the *Myo15-Cre* conditional mouse model and alleviated vestibular deficits in the constitutive knockout mouse model of USH1F. a Dual AAV vectors together encoding PCDH15-CD2 were injected via the round window membrane (RWM) into either *Myo15-Cre* conditional knockout mice or constitutive knockout *Pcdh15^-/-^* mice at age P1, at a dose of 5×10^10^ VGC. At 5 weeks, treated animals were assayed for the rescue of hearing and vestibular function, and tissue processed for histology. **b** Average tone ABR thresholds as a function of frequency and click thresholds for 5 weeks uninjected *Pcdh15^fl/fl^* hearing control mice (*n* = 9), uninjected *Pcdh15^fl/fl^*, *Myo15-Cre ^+/-^* conditional knockout mice (*n* = 10), and conditional knockout mice treated with dual-AAV (n = 23). The dark red ABR line is the average of the 23 individual treated ABRs. In bright red, the average of the three best performers is displayed. **c** Average distortion-product otoacoustic emission (DPOAE) thresholds and average DPOAE emission amplitudes at 16 kHz, for 5 weeks hearing control mice (*n* = 9) and uninjected *Pcdh15^fl/fl^*,*Myo15-Cre ^+/-^* mice (*n* = 10), and for those injected with dual-AAV (*n* = 23). **d** Representative open field path outlines of wild-type C57BL/6 mice, uninjected *Pcdh15^fl/fl^*, *Myo15-Cre ^+/-^* conditional knockout mice, homozygous *Pcdh15^-/-^* knockout mice and *Pcdh15^-/-^* knockouts treated with dual AAVs. **e** Summary of total rotations in wild-type C57BL/6 *(n*=5), uninjected *Pcdh15^fl/fl^*, *Myo15-Cre ^+/-^* late conditional knockout mice *(n*=5), homozygous *Pcdh15^-/-^* constitutive knockout mice *(n*=4) and *Pcdh15^-/-^* treated with dual AAVs *(n*=13). Untreated *Pcdh15^-/-^* mice exhibited severe circling behavior, while in dual-AAV treated animals vestibular function was fully restored to normal level. **f** wild-type C57BL/6 *(n* =5) and *Pcdh15 ^fl/fl^,Myo15-Cre ^+/-^* (n=5) mice 5 weeks old swim well, keeping their heads above water for more than 60 s. *Pcdh15^-/-^* mice *(n*= 4) were unable to swim normally and were quickly removed. The dual-AAV delivery of PCDH15 successfully restored swimming function in *Pcdh15^-/-^* mice (*n*=12). g *Pcdh15^-/-^* mice showed much shorter latency to fall on the rotarod test *(n*= 4), while treatment with dual-AAV restored rotarod performance to wild-type level (*n*=12). All data are presented as mean ± SEM. Source data are provided as a Source Data file.

We first assessed hearing in treated mice by recording ABRs to broadband clicks and tone bursts (**Fig. 2b**). We found that *Pcdh15^fl/fl^, Myo15-Cre ^+/-^* injected at P1 with dual AAVs encoding either PCDH15 or HA-PCDH15 showed robust rescue of hearing (**Fig. 2b**). The thresholds in rescued animals treated with PCDH15 either with or without the HA tag were the same (**Supplementary Fig. 1a**), confirming that HA-tagged PCDH15 is fully functional. We therefore combined the HA-tagged and non-tagged mice in **Fig. 2b,c**. In some mice, hearing was near that of untreated *Pcdh15^fl/fl^* control mice lacking Cre. Untreated *Pcdh15^fl/fl^,Myo15-Cre ^+/-^* mice were deaf at P30 as tested by ABR, with thresholds above 85 dB, the highest level tested. *Pcdh15^fl/fl^* normal hearing control mice injected with dual AAVs had normal thresholds indicating no sign of toxicity for hearing (**Supplementary Fig. 1b**).

DPOAE measurements also demonstrated strong rescue in dual-AAV treated mice (**Fig. 2c**), with thresholds near those of hearing control *Pcdh15^fl/fl^* Cre^-^ mice. The three best-performing mice had thresholds equal to controls. Similarly, DPOAE amplitudes measured at a representative midrange frequency (16 kHz) showed rescue near the hearing control level in treated *Pcdh15^fl/fl^, Myo15-Cre ^+/-^* conditional knockout mice. We observed that *Pcdh15 ^fl/fl^* control mice injected with the dual AAVs exhibited normal DPOAE thresholds, further indicating absence of toxicity. These experiments use a late-deletion mouse model which allows normal hair-cell development before gene deletion, but the model may not represent the phenotype of the many Usher 1F patients who lack PCDH15 at all ages. The experiments nevertheless show that dual-AAV delivery produces a functional PCDH15 and restores hearing.

Surprisingly, *Pcdh15^fl/fl^,Myo15-Cre ^+/-^* mice did not show a behavioral vestibular deficit, assessed by swimming, open-field locomotion and rotarod tests (**Fig. 2d-g**). We previously speculated that the presence of functional PCDH15 in the first postnatal week allowed proprioceptive input to be linked with vestibular sensation, so that the mice could adequately perform locomotory tasks even after deletion of PCDH15^28^. To test efficiency of dual-AAV delivery of PCDH15 in rescue of vestibular function in a more severe model, we used a constitutive knockout mouse model bearing the R245X truncation mutation (*Pcdh15^R^*^245^*^X/R^*^245^*^X^*, here referred to as *Pcdh15^-/-^*)^28^. These mice exhibited a severe vestibular phenotype including intensive head bobbing, circling behavior, hyperactivity, difficulty swimming, and inability to remain on a turning rotarod^28^ (**Fig. 2d-g**). AAV vectors were administered via the RWM into *Pcdh15^-/-^* mice at P1, with a dose of 1.0×10^11^ VGC. In mice 5 weeks of age, we assessed vestibular function rescue with swimming, open field locomotion and rotarod tests. In all three tests, the dual-AAV delivery of full-length PCDH15-CD2 restored function to wild-type levels (**Fig. 2d-g**).

### Dual-AAV delivery of PCDH15-CD2 restored stereocilia bundle morphology, tip links, and mechanotransduction in the Pcdh15^fl/fl^, Myo15-Cre conditional knockout mouse

Next, we assessed the rescue of stereocilia bundle morphology and mechanotransduction using dual-AAV delivery, employing both a fluorescent phalloidin actin label and scanning electron microscopy. Actin labeling of hair bundles revealed that at 5 weeks, untreated conditional knockout mice exhibited disorganized bundles or had lost their hair bundles entirely (**Fig. 3a,b**). This disorganization included the loss of short and middle row stereocilia, with only some of the tall row stereocilia remaining. In contrast, dual-AAV delivery of PCDH15 led to a robust rescue of hair cell morphology, with bundles at P35 closely resembling those in *Pcdh15^fl/fl^* Cre-control mice (**Fig. 3a,b**).

**Fig. 3.**
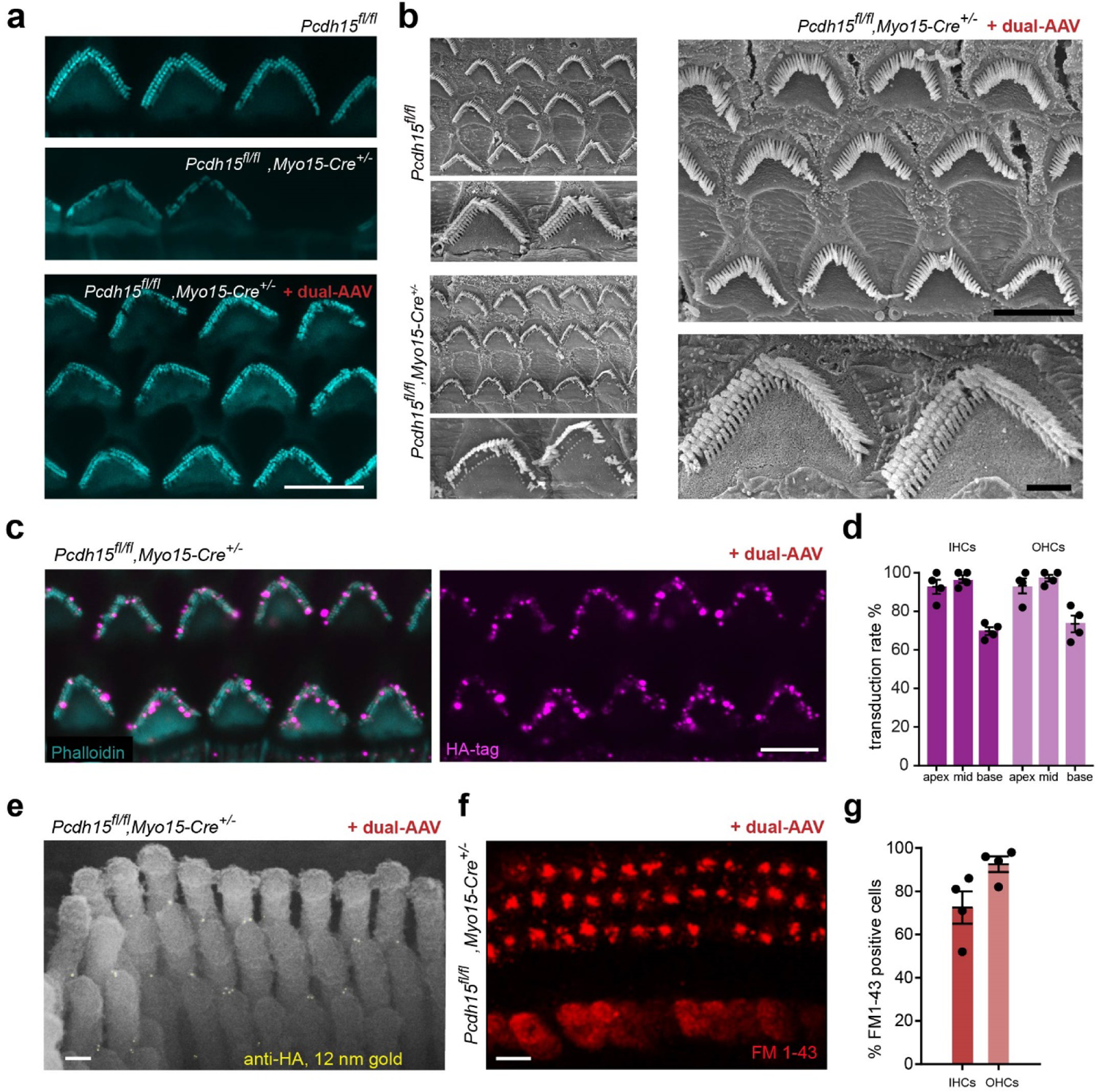
Delivery of dual-AAV restores the morphology of stereocilia bundles, mechanotransduction, and tip-links in the Myo15-Cre conditional knockout mouse model. a Representative confocal microscopy images captured from the middle turn of 5-week-old cochleas, showing outer hair cells (OHCs) stained with phalloidin. The panels illustrate OHCs from *Pcdh15^fl/fl^* hearing control mice, uninjected Myo15-Cre conditional knockout mice, and knockout mice that received dual-AAV injections. b Scanning electron micrographs of hair bundles of a hearing control mouse and untreated conditional knockouts. In the knockouts, the bundles are severely disrupted. However, in the OHCs of a conditional knockout treated with dual-AAV, normal bundle morphology was restored. c Representative confocal microscopy images at 5 weeks from middle turn of the cochlea show anti-HA staining at the tips of stereocilia in knockout cochleas treated with dual-AAV encoding HA-PCDH15 at P1. d Transduction efficiency in IHCs and OHCs at 5 weeks in treated *Pcdh15^fl/fl^,Myo15-Cre ^+/-^* conditional knockout mice (*n* = 4). Data are presented as mean± SEM. e Immunogold scanning electron microscopy localization of HA-tagged PCDH15 in OHC stereocilia of treated KO mice. Multiple 12-nm gold beads (light yellow) were detected in a scanning electron microscopy image, confirming that HA-tagged PCDH15 goes to the tips of stereocilia, except the tallest. f Rescue of FM1-43 uptake in a treated cochlea. g Average percentage of IHCs and OHCs loaded with FM1-43 in conditional knockout mice injected with dual-AAV (*n* = 4). Data are presented as mean values ± SEM.

The successful restoration of hearing and bundle morphology further indicates that PCDH15-CD2 is effectively expressed and accurately targeted within the hair cells. To assess the proper trafficking and localization of exogenous PCDH15 at the tips of cochlear stereocilia, we utilized the AAV-HA.PCDH15 5’ + AAV-PCDH15 3’ vector pair. The AAV-HA.PCDH15 5’ vector incorporates a hemagglutinin (HA) tag at the N-terminus of PCDH15. Immunofluorescence imaging confirmed that conditional knockout mice injected at P1 with AAV-HA.PCDH15 5’ + AAV-PCDH15 3’ exhibited clear immunoreactivity to the HA tag. Importantly, this labeling was accurately located at the tips of stereocilia (**Fig. 3c**). Hair-cell expression of HA-PCDH15 varied from apex to base, with 93–98% of apical and middle IHCs and OHCs transduced and 70–74% of basal hair cells transduced (**Fig. 3d**).

With scanning electron microscopy and a secondary antibody conjugated to 12 nm gold beads, we observed the location of HA.PCDH15 within hair bundles. Gold beads were specifically localized on the tips of short and middle row stereocilia, precisely at the position of the tip links (**Fig. 3e**). This confirms the accurate trafficking and localization of HA-PCDH15 at age 5 weeks.

In order to evaluate the rescue of mechanotransduction in hair cells with dual-AAV delivery, we dissected apical and mid-apical regions of the cochleas at 5 weeks. We then briefly applied FM1-43 dye directly onto the exposed epithelium, and subsequently, a solution of SCAS. We quantified the proportion of FM1-43-positive cells in the cochlea. FM1-43 labeling revealed a restoration of mechanotransduction in approximately 93% of outer hair cells (OHCs) and 73% of inner hair cells (IHCs) (**Fig. 3g**).

### PCDH15 gene structure and disease-associated variants

*Pcdh15* is expressed as one of three primary transcripts, namely the CD-1, CD-2, and CD-3 isoforms. These isoforms primarily differ in the 3’ end of the gene. The CD2 splice form is necessary for PCDH15 function in hair cells ^17,38^, though it remains uncertain which isoform is predominant in the retina. In order to investigate this we analyzed *PCDH15* exon structure and disease-associated variants.

Disease-associated variants were systematically collected from the Deafness Variation Database (PMID: 30245029), accessed in August 2021. These variants were then mapped to their respective positions on the human NM_001142769 transcript and color-coded based on the associated disease phenotype: variants associated with non-syndromic hearing loss (DFNB23), variants linked to USH1F with both deafness and blindness, and variants connected with non-syndromic retinitis pigmentosa (NSRP). It is noteworthy that a specific variant, p.P1796fs, impacts the protein-coding sequence exclusively within the CD-1 isoform. This variant has been documented as a causative factor for NSRP (PMID: 27208204), suggesting a fundamental role of the CD-1 isoform in retinal function (**Fig. 4a**).

**Fig. 4.**
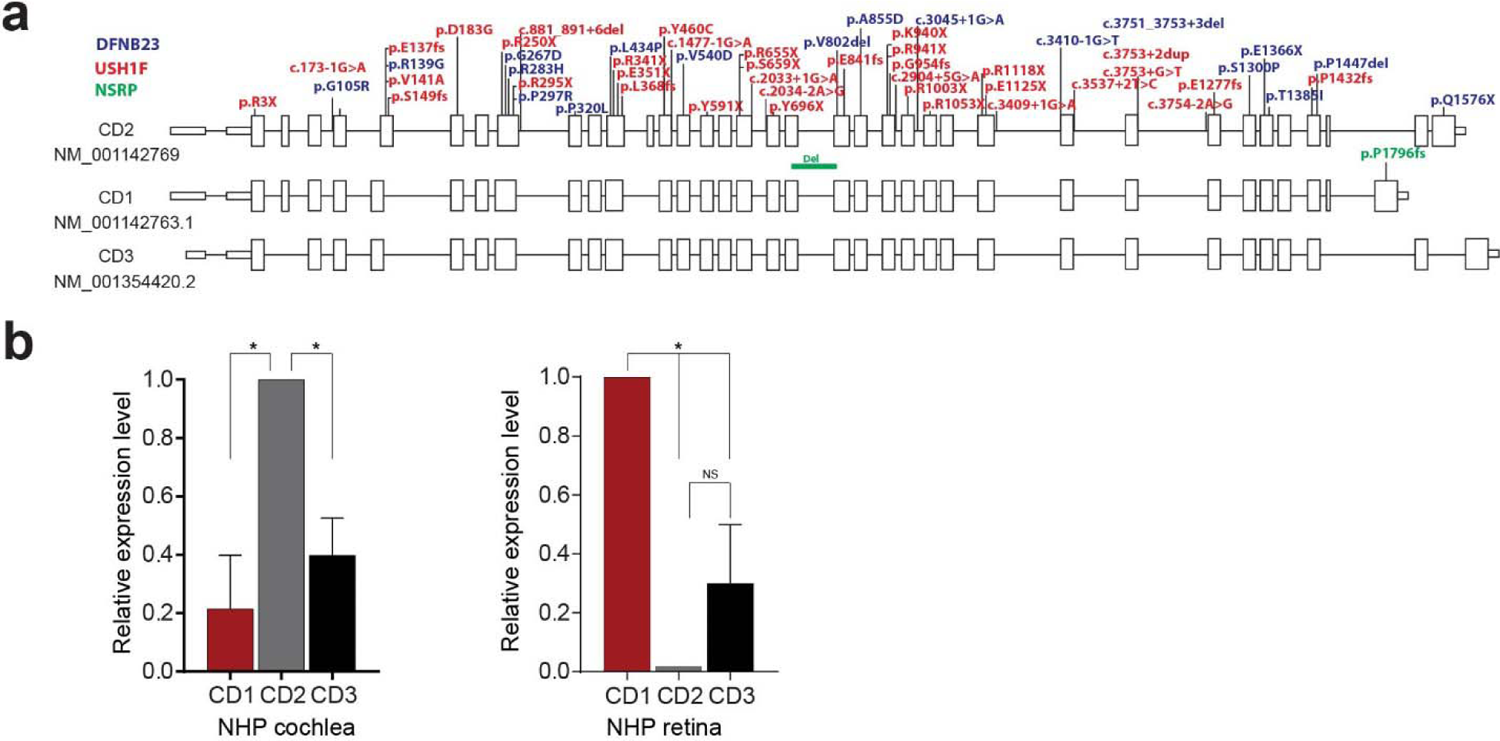
*PCDH15* exon structure and disease-associated variants. **a** Gene schematic of PCDH15-CD2 and reported disease-causing variants. Three primary transcripts of PCDH15 translate three isoforms (CD-1, CD-2, and CD-3), which primarily differ in the 3’ end of the gene. Disease variants were gathered from the Deafness Variation Database (PMID: 30245029), accessed Aug 2021. Disease variants are plotted based on their location on transcript NM_001142769 and color-coded based on associated disease phenotype; blue, non-syndromic hearing loss (DFNB23), red, Usher Syndrome Type 1F (USH1F), or green, non-syndromic retinitis pigmentosa (NSRP). Note that the variant p.P1796fs impacts the protein-coding sequence of only the CD-1 isoform and has been reported to cause NSRP (PMID: 27208204), suggesting the CD-1 is required for retinal function but not hearing. Similarly, p.Q1576X in CD2 affects hearing but not vision. **b** qPCR evaluation of PCDH15-CD1, PCDH15-CD2 and PCDH15-CD3 expression in primate cochlea and retina. Data are presented as mean ± SEM. Source data are provided as a Source Data file.

We used RT-qPCR on an inner ear and eye of non-human primates (NHPs) to reveal tissue-specific PCDH15 isoform expression. In the NHP inner ear, the predominant splice form of the C-terminal region was identified as CD2, with some detectable expression of CD3 (**Fig. 4b**). Within the NHP retina, CD1 expression was far higher than that of CD2; again, there was some CD3 expression. These findings highlight the tissue-specific variation in PCDH15 isoform expression, with CD2 being prominent in the inner ear and CD1 prevailing in the NHP retina.

### Evaluation of dual-AAV in human retina organoids

Retinal degeneration in USH 1F patients occurs over decades providing a window of opportunity for the treatment of retinal degeneration. Gene therapy could target degenerating photoreceptors in the retina to halt visual field loss, but it requires testing of a therapeutic construct in a model that is relevant for the human retina. We derived retinal organoids from induced pluripotent stem cells and tested gene delivery using the dual-AAV vector strategy (**Fig. 5a**). First, we examined photoreceptor ultrastructure within retinal organoids and the localization of PCDH15 within photoreceptors. Immunofluorescence imaging of cryosections labeled with anti-PCDH15 antibodies revealed distinct staining in the photoreceptors, especially at the distal ends of the inner segments (**Fig. 5b**). Additionally, scanning electron microscopy analysis revealed that the majority of photoreceptors exhibited well-developed inner segments, while a subset displayed outer segments and connecting cilia. Notably, we observed the emergence of nascent calyceal processes at the apical ends of the inner segments (**Fig. 5c**). Further investigation using immunogold scanning electron microscopy, where human retinal organoids were subjected to immunostaining with anti-PCDH15 primary antibodies and subsequent labeling with 12-nm gold-conjugated secondary antibodies, demonstrated the trafficking of PCDH15 to the surfaces of the inner segments and the nascent calyceal processes of the photoreceptors (**Fig. 5e**).

**Fig. 5.**
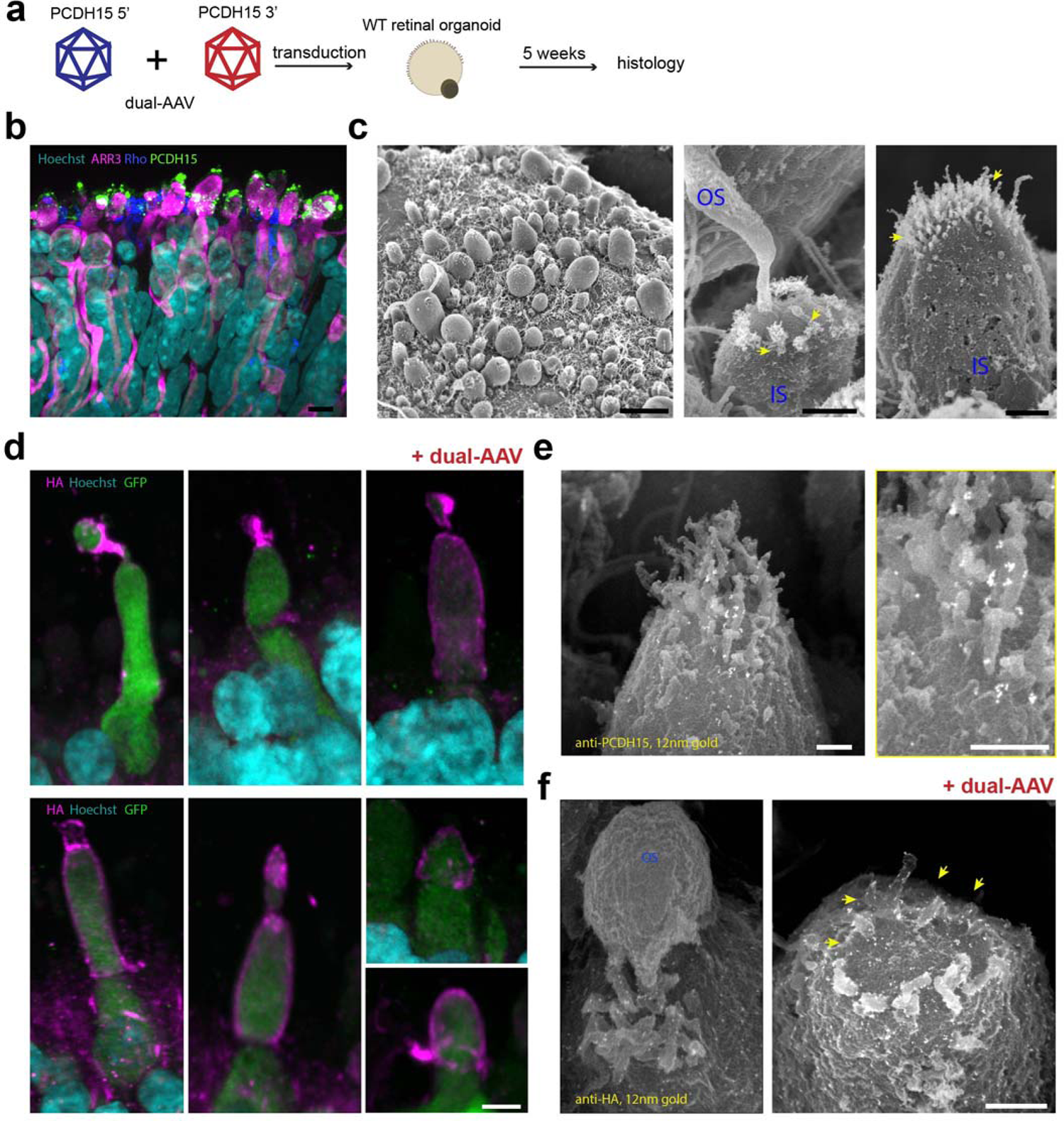
Expression of PCDH15 in human retinal organoids by dual-AAV vectors. **a** Organoids were transduced with dual-AAV vectors and five weeks later expression was analyzed by antibodies to the HA tag. **b** Representative confocal microscopy images captured from cryosections of non-transduced human retina organoids. Anti-PCDH15 labeling was detected in the photoreceptors, especially at the distal ends of outer segments. **c** Scanning electron microscopy of human retinal organoids. The finger-like calyceal processes (yellow arrows) protrude from the apical region of the inner segments (IS) of both rod (middle) and cone (right) photoreceptors. **d** Because the CMV promoter is only present on the 5’ vector, HA.PCDH15 and GFP were only expressed when both the 5’ and 3’ vectors were added to the organoids. The HA.PCDH15 is shown as magenta and GFP as green. In these images, HA.PCDH15 localizes to the junction of inner and outer segments of cone photoreceptors. **e** Immunogold scanning electron microscopy labeling of non-transduced human retinal organoids immunostained with anti-PCDH15 primary antibody and 12 nm gold-conjugated secondary antibody. PCDH15 trafficked to the surface of inner segment and to nascent calyceal processes of photoreceptors. **e** Immuno-gold scanning electron microscopy confirmed localization of HA-tagged hsPCDH15 to calyceal processes and IS of human retinal organoids transduced with dual-AAV. Multiple 12-nm gold beads were detected. Scale bars: **b, d** 5 µm, **c** 10 µm, **e, f** 0.5 µm.

Next, we packaged CMV-GFP into an AAV9-PHP.B capsid. Human retinal organoids were incubated with 8.5×10^11^ VGC and five weeks later evaluated for GFP expression (**Fig. 5a**). We found that AAV9-PHP.B robustly transduced photoreceptors of retinal organoids (**Supplementary Fig. 2a**).

For gene expression studies in retina, we used the human PCDH15-CD1 isoform (NM_001142763.1) tagged with HA. We engineered dual-AAV hybrid vectors so that each vector encoded part of the full-length protein. As for mouse vectors, Vector 1 (AAV-HA.PCDH15 5’) included a CMV promoter, the N-terminal half of the *PCDH15* coding sequence with HA tag, and a splice donor site followed by the HR recombinogenic sequence. Vector 2 (AAV-PCDH15 3’) included the HR sequence, a splice acceptor, the C-terminal half of the *PCDH15* coding sequence, and the GFP coding sequence following an independent ribosomal entry site (IRES). We transduced retinal organoids derived from human induced pluripotent stem cells using these dual AAVs, separately or together. After five weeks, we assessed the expression of GFP and HA-tagged PCDH15 employing immunofluorescence and immunogold scanning electron microscopy imaging. Because the organoids presumably expressed the endogenous human PCDH15, we detected the AAV-delivered PCDH15 with antibodies to HA rather than to PCDH15 itself.

Notably, we did not detect the presence of the HA tag when organoids were transduced with either AAV-HA. PCDH15 5’ or AAV-PCDH15 3’ alone (**Supplementary fig. 2b, 2c**). However, anti-HA antibody labeling revealed expression of HA-PCDH15 in organoids transduced with the dual AAVs, and further showed that the HA tag was specifically localized on the surfaces of the inner segments and at the junction between the inner and outer segments, precisely where the development of calyceal processes occurs. This observation confirmed that the dual-AAV approach, initially developed for the inner ear, successfully transduced and expressed full-length PCDH15 in photoreceptors, with specific localization to the calyceal processes and inner segments of photoreceptors (**Fig. 4d and 4f**).

### Evaluation of dual-AAV delivery in NHP retina

Finally, we assessed the efficiency of dual-AAV in an old-world non-human primate (NHP), which represents the most pertinent animal model for transgene delivery to the human eye. In the green monkey (*Chlorocebus sabaeus*), we administered dual-AAV through subretinal injection, a well-established and effective route for delivering therapeutic agents to the retina. Specifically, in one eye AAV injection produced three blebs in the superior, inferior, and temporal regions of the retina, each containing dual-AAV at a dosage of 2.5 x 10^12^ VCG per bleb (totaling 7.5 x 10^12^ VCG per eye), with each bleb containing 50 µl of virus. The second eye served as a control and received an injection of a formulation buffer consisting of PBS/0.001% F-68, with the same distribution of three blebs in the superior, inferior, and temporal regions of the retina, each containing 50 µl. Before the injection procedure, the monkey underwent a comprehensive baseline evaluation. The general assessment included the determination of AAV9 neutralizing antibody (nAb) seronegativity, an evaluation of general well-being, and a thorough examination of ocular health. To verify the successful formation of subretinal blebs, we employed OCT immediately following the surgical procedure. In general, the procedure was well tolerated by the animal, and it was closely monitored for a duration of 9 weeks post-injection. Subsequently, we conducted an analysis of retinal tissues using histological methods (**Fig. 6a**).

**Fig. 6.**
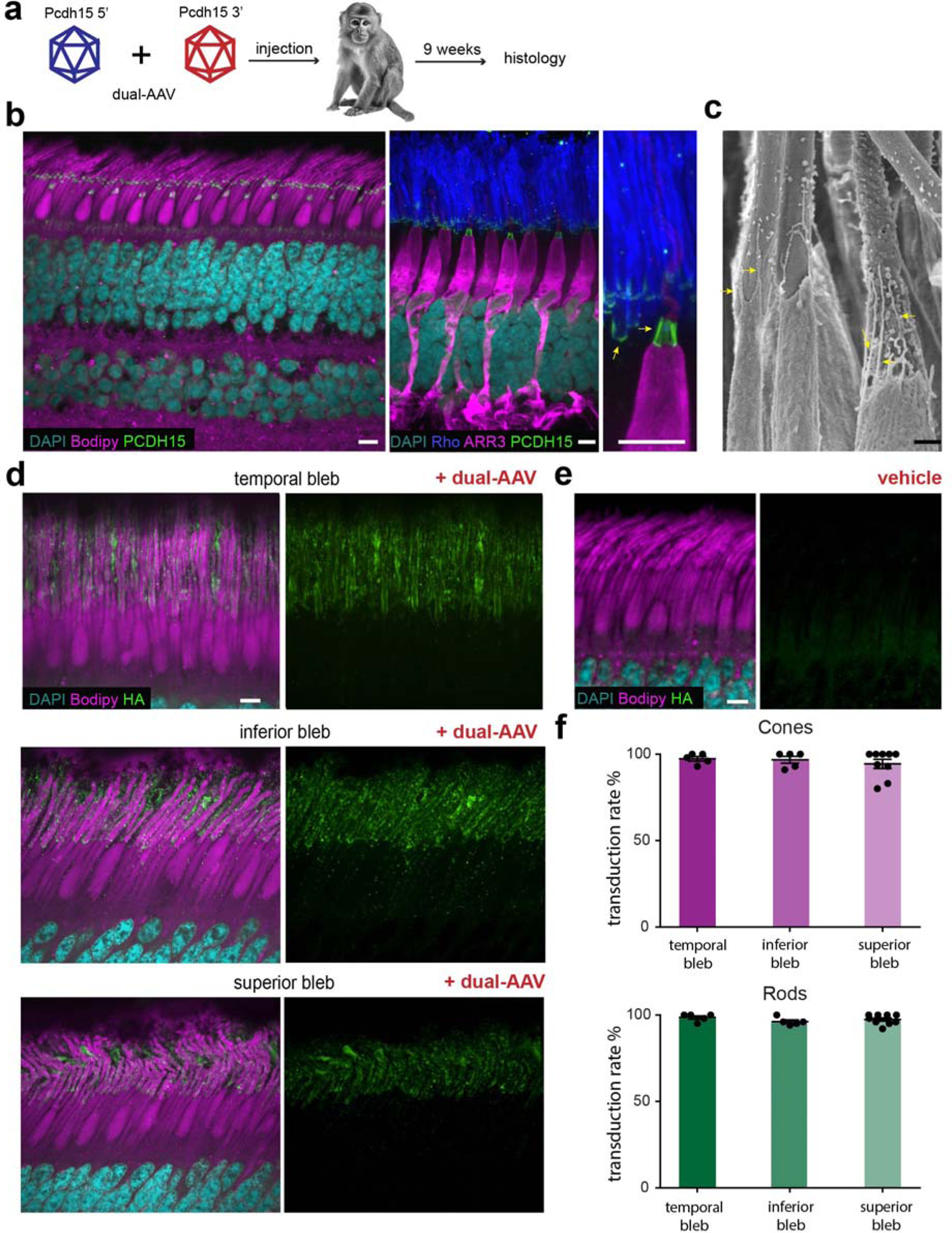
Localization of endogenous and dual-AAV-delivered PCDH15 in nonhuman primate retina. **a** Dual AAV vectors were injected subretinally into an NHP eye. At 9 weeks, the treated retina was assayed for expression and localization by histology. **b** Immunofluorescence labeling of endogenous PCDH15 (green) in a vehicle-injected control retina. An antibody to cone arrestin marks cone photoreceptors (magenta) and an antibody to rhodopsin marks outer segments of rods (blue). PCDH15 is located at the junction between inner and outer segments and along the calyceal processes in both cones and rods. **c** Scanning electron micrograph of the inner–outer segment interface in a cone photoreceptor (right) and rod photoreceptors (left). The calyceal processes (arrows), protrude from the apical inner segment of photoreceptors to surround the outer segment. **d** Representative confocal microscopy images captured from cryosections of temporal, superior and inferior blebs created by the dual-AAV injection. An anti-HA signal was detected in the photoreceptors injected with dual AAVs but not in the eye injected with vehicle (**e**). **f** Transduction efficiency in cone and rod photoreceptors injected with dual AAVs. Scale bars: **b, d, e** 5 µm; **c** 1 µm. Data are presented as mean ± SEM. Source data are provided as a Source Data file.

We initially determined the normal localization of PCDH15 in photoreceptors of the control eye using an anti-PCDH15 antibody and the fluorescent dye BODIPY, which labels lipid membranes including those in cones and rods. Our findings confirmed that PCDH15 localizes, as previously demonstrated, to the interface between the inner and outer segments of both rods and cones (**Fig. 6b**). Next, we conducted a more detailed examination of the outer-inner junction by immunolabeling the retina with antibodies specific to cone arrestin and rhodopsin. The cone arrestin counterstain provided insights into the comprehensive cone anatomy, revealing well-preserved structures that closely matched those observed in scanning electron microscopy images (**Fig. 6c**). PCDH15 staining formed a barrel-like structure surrounding the base of the outer segment (**Fig. 6b**), which correspond to the calyceal processes seen in scanning electron microscopy. In contrast, rhodopsin labeled the outer segment of rod cells, where PCDH15 was observed as a thin layer at the base of the outer segment, with a few extensions along the outer segment (**Fig. 6b**). This pattern correlates with the fewer number of calyceal processes in rods compared to cones (**Fig. 6c**).

Subsequently, we examined samples obtained from the eye injected with dual AAVs encoding HA.PCDH15. Immunostaining against the HA-tag demonstrated robust overexpression of PCDH15 within the photoreceptor layer in the inner/outer segment junction, which extended along outer segments of cones and rods (**Fig. 6d**). We quantified the transduction efficiency in all three regions. Anti-HA labeling demonstrated high numbers of transduced cones and rods throughout bleb areas with ∼94–97% of cones labeled and ∼96–98% of rods. No labeling was detected in the eye injected with vehicle (**Fig. 6e**).

Altogether, the results show that the dual-AAV approach, effective in human retina organoids, was successfully translated to NHPs. High levels of exogenous full-length PCDH15 were reached in the macaque eye after subretinal injection (**Fig. 6f**).

## Discussion

Significant progress has been achieved recently in the field of gene therapy for deafness ^45-48^ and blindness ^14,26,49–51^. This progress has usually involved gene-addition therapy, which has proven effective for small genes with coding sequence that can be accommodated within a single AAV vector (<4.7kb). However certain genes, including many responsible for Usher syndrome, have coding sequences that exceed the capacity of AAV vectors ^1,2,52^.

Here, we demonstrate the successful application of a dual-AAV vector strategy for gene augmentation therapy in audio-vestibular and visual sensory systems using four different animal models: constitutive and conditional mouse models of USH1F deafness and balance dysfunction, cultured human retinal organoids and intact nonhuman primate retina.

We employed a hybrid dual-AAV approach ^31–35,37^ to deliver the full-length PCDH15 coding sequence, which is too large to fit within a single AAV capsid, by using two AAV vectors (AAV-Pcdh15 5’ and AAV-Pcdh15 3’) encoding different portions of the Pcdh15 coding sequence. Upon transduction of target cells with both vectors, a highly recombinogenic AK sequence facilitated recombination of DNA from the two vectors, resulting in the production of full-length PCDH15 mRNA and protein.

In this study, we selected the AAV9-PHP.B capsid for delivery and demonstrated its excellent ability to transduce sensory cells within the mouse cochlea^41–43^, as well as photoreceptors in retinal organoids and in non-human primate retinas.

The dual-AAV approach was initially validated *in vitro* using HEK 293 cells, confirming efficient recombination and splicing that led to PCDH15 protein production. *In vivo* experiments with conditional knockout *Pcdh15^fl/fl^, Myo15-Cre* mice, which suffered profound hearing loss if left untreated, showed that dual-AAV delivery of PCDH15-CD2 ^17,38^ robustly restored auditory function, with some mice reaching control-level thresholds. This treatment also restored stereocilia bundle morphology and mechanotransduction in hair cells. Immunofluorescence and immunogold scanning electron microscopy confirmed the precise trafficking and localization of exogenous PCDH15 at cochlear stereocilia tips. Treated control mice showed no signs of hearing-related toxicity. Vestibular function was also assessed using a constitutive knockout *Pcdh15-/-* mouse model, which exhibited severe vestibular dysfunction. The dual-AAV treatment restored vestibular function to near-normal levels in these mice, as indicated by improved performance in various behavioral tests.

The utilization of human retinal organoids derived from human induced pluripotent stem cells has enabled the examination of both retinal development and the study of retinal diseases ^53–58^. They consist of three nuclear layers and two synaptic layers, replicating the cellular organization observed in the adult retina, and that they exhibit photoreceptor responsiveness to light stimuli and facilitate synaptic transmission of visual information, resulting in light responses in second or third order retinal cells ^56^. We therefore examined the photoreceptor ultrastructure within retinal organoids and the localization of PCDH15. The emergence of nascent calyceal processes at the apical ends of the inner segments, where we found PCDH15, validates the suitability of organoids as a model for PCDH15 delivery. Immunofluorescence and immunogold scanning electron microscopy imaging confirmed the successful transduction and localization of HA-tagged PCDH15 in photoreceptors, particularly in the calyceal processes and inner segments.

Old world monkeys are likely the most suitable model for investigating ocular diseases. Since only primates possess a macula, NHP models play a crucial role not just in uncovering the biological mechanisms behind high-acuity vision but also in advancing therapeutic development ^59^. Recent years have seen successful transitions from basic research to clinical trials and even the approval of the first treatments for inherited and age-related retinal dystrophies ^60^. In our study subretinal injection of dual-AAV in an NHP demonstrated high levels of exogenous PCDH15 expression in both rods and cones, indicating the potential applicability of this approach to human patients.

In summary, the dual-AAV approach holds significant promise for the development of gene replacement therapy for USH1F. The potential restoration of both auditory and vestibular functions indicates that the dual-AAV strategy could present a promising therapeutic option, especially when considering that treatments for deafness, such as cochlear implants, are already available, whereas there are currently no treatments for vestibular dysfunction which can be offered for patients with USH1F. Furthermore, the successful application of the dual-AAV approach in human retinal organoids and NHPs provides a foundation for future translational studies aimed at treating retinal degeneration associated with USH1F. These findings provide a strong foundation for further research and the development of potential treatments for Usher syndrome type 1F and related conditions.

## Methods

### Study approval

All animal studies were conducted in compliance with ethical regulations according to protocol IS00001452 approved by the Institutional Animal Care and Use Committee (IACUC) at Harvard Medical School, Boston, and were performed according to the NIH guidelines.

### Dual-PCDH15 study design

The dual-PCDH15 strategy was designed based on the hybrid strategy^30,31,33^. To promote recombination between the 5’ and 3’ viruses, we inserted the highly recombinogenic (AK) sequence derived from the F1 phage genome ^32^. The 5’ virus harbors SD site upstream of the AK sequence, while the 3’ virus harbors SA site immediately downstream of the AK sequence. The coding sequence of the mouse CD2-1 isoform of PCDH15 was used for the initial design (NM_001142742.1). Experiments in retinal organoids and in the green monkey retina used the coding sequence for the human CD1-1 isoform (NM_001142763.2). The AAV expression cassettes tested in this study are shown in **Fig. 1a**. We used an AAV transgene plasmid, flanked by AAV2 ITRs. For all experiments, we utilized a 584-bp CMV promoter, which we had previously demonstrated to be effective in the cochlea^27^. Additionally, 3’ viruses were engineered to include an IRES GFP element to facilitate the co-translation of a fluorescent marker. These constructs also incorporated the woodchuck hepatitis virus post-transcriptional regulatory element (WPRE) and a bovine growth hormone (BGH) poly(A) sequence.

### Mouse models

Animal handling, breeding, and all procedures on mice were performed in compliance with NIH ethics guidelines and under a protocol approved by the Animal Care Committee of Harvard Medical School. Mice were housed and bred in the Harvard Medical School animal facility. All studies were performed on *Pcdh15^fl/fl^,Myo15-Cre* and *Pcdh15*^R245X/ R245X^ mice, which were previously described^27,28^.

### Viral vector production

Most AAVs produced in this study were manufactured by the Viral Vector Core at Boston Children’s Hospital. For NHP experiments, AAVs were produced by PackGene. Specifically, AAV9-PHP.B vectors were generated through the use of HEK 293 cells. This was achieved by polyethylenimine-mediated co-transfection of pAAV transfer plasmid, pHelper plasmid, and RepCap plasmid pUCmini-iCAP-PHP.B. After 120 hr post-transfection, both the media and cells were collected. Subsequently, AAV9-PHP.B viruses were extracted and subjected to ultracentrifugation using a discontinuous density iodixanol (OptiPrep, Axis-Shield) gradient. Following ultracentrifugation, AAV vector-containing iodixanol fractions were isolated and concentrated via diafiltration. The purified AAV vectors were quantified using q-PCR, divided into single-use aliquots, and stored at −80°C until required, with thawing taking place immediately before in vivo injections.

### Transduction of HEK 293 cells

HEK 293 cells (ATCC, #CRL-1573) were plated on glass coverslips in DMEM with 10% FBS (Gibco) and penicillin/streptomycin (Pen/Strep; Invitrogen). On the following day, cells were transfected with Lipofectamine 3000 according to the manufacturer’s protocol. Plates were incubated at 37 °C for 24 hr and then at 30 °C for an additional 48 hrs to promote high levels of protein expression.

To transduce HEK 293 cells with AAV9-PHP.B, the cells were transduced when they reached a low confluency of approximately 40%, using a multiplicity of infection of approximately 7 × 10^5^. The viruses were diluted in 200 µl of DMEM containing 1% FBS, penicillin, and streptomycin, and then applied directly onto coverslips to create a hydration bubble. These coverslips were then placed in a 37°C incubator. After 16-24 hours, the cells were washed with media containing 1% FBS and 0.5x Pen/Strep and subsequently replenished with 2 mL of fresh media. They were then cultured at 30 °C for an additional 48 hrs to promote high levels of protein expression. Cells were collected for mRNA or fixed for immunofluorescence.

### RNA extraction, cDNA production, reverse transcription, PCR amplification, and sequencing

Total RNA was extracted from transduced HEK 293 cells using the Zymo Quick-RNA Microprep Kit (Zymo Research, #R1050). Reverse transcription was performed with Invitrogen’s SuperScript IV VILO Master Mix with ezDNAse enzyme (Invitrogen, #11766050). PCDH15 cDNA was amplified with a forward primer which anneals to sequence in the 5’ virus (5’-aggatgaaaacgatcaccccc-3’) and a reverse primer which anneals to sequence in the 3’ virus (5’-ggtatgatgagccggtaggc-3’). The expected product size for properly spliced PCDH15 cDNA splice junction site is 154 bp. The DNA Gel Extraction Kit (Monarch, #T1020S) was used to gel-extract PCR products. Purified PCR products were subcloned, transformed into competent cells, mini-prepped, and Sanger sequenced using the NEB PCR Cloning Kit (NEB, #E1202S).

### Expression in human retinal organoids

Wild-type human retina organoids were generated and maintained as was described previously ^56^. These organoids were subsequently transduced with AAV vectors in a 96-well plate using the following dosages: dual-AAV (AAV-HA.PCDH15 5’ + AAV-PCDH15 3’), 1.7×10^12^ VGC; AAV-HA.PCDH15 5’, 8.5 ×10^11^ VGC; AAV-PCDH15 3’, 8.5 ×10^11^ VGC; AAV-CMV-GFP, 8.5 ×10^11^ VGC. Transduced organoids were maintained in 50 µL of 3:1 N2 medium at 37°C in 5% CO2. Composition 3:1 N2 as published in Cowan et al. 2020: DMEM (Gibco, #10569-010) supplemented with 20% Ham’s F12 Nutient mix (Gibco, #31765-027), 10% heat-inactivated fetal bovine serum (Millipore, #ES-009-b), 1% N2 Supplement (Gibco, #17502-048), 1% NEAA Solution (Sigma, #M7145), 100uM taurine (Sigma, #T0625) and I uM retinoic acid (Sigma, #R2625).

After 4 hr, 50 µL of media was added to each well. One day later, 100 µL of media was added to each well. After 24 hr and every 48 hr thereafter, the solution was completely exchanged with fresh media. Five weeks later, samples were fixed with 4% formaldehyde.

### Expression in nonhuman primate retina

For in vivo delivery, we chose an old-world primate, the green monkey (*Chlorocebus sabaeus*) for greater relevance to human. Injections were carried out by Virscio (New Haven, CT). Before injection, the animal underwent a thorough ophthalmological assessment conducted by a veterinary ophthalmologist, and baseline screening to assess AAV9 neutralizing antibody (nAb) seronegativity. The animal received methylprednisolone (40 mg IM) weekly for 4 weeks, starting on the day prior to dosing. Anesthesia was achieved with intramuscular ketamine (8 mg/kg) and xylazine (1.6 mg/kg) to effect, and pupil dilation with topical 10% phenylephrine, 1% tropicamide and/or 1% cyclopentolate. After placing the scleral ports, a contact vitrectomy lens was positioned on the cornea using 0.9% saline as a coupling agent. A 25-gauge light pipe was inserted through the left scleral port into the vitreous cavity for intraocular illumination. Simultaneously, a subretinal cannula was introduced through the second scleral port. The cannula was gently advanced to touch the retinal surface in a specific location. Once the retinal surface showed a slight blanching at the point of contact, the vector was administered through the cannula. Three blebs were introduced, in the superior, inferior, and temporal regions of the retina, each containing dual-AAV at a dosage of 2.5 x 10^12^ VGC per bleb (totaling 7.5 x 10^12^ per eye), with each bleb receiving 50 µl of virus. The second eye served as a control and received an injection of a formulation buffer consisting of PBS with 0.001% F-68, with the same distribution of three blebs in the superior, inferior, and temporal regions of the retina, each containing 50 µl.

### RNA extraction, cDNA production, reverse transcription-quantitative polymerase chain reaction (RT-qPCR) in NHP

Three cynomolgus monkey (*Macaca fascicularis*) cadavers were acquired post-euthanasia from an unrelated study at Massachusetts General Hospital and transferred to Harvard Medical School as approved by IRB Ex Vivo Animal Tissue Importation Amendment ID 14-111-A02. Inner ear tissue and retinas were extracted and transferred into Trizol solution surrounded by a liquid nitrogen bath. Once cochlear and retinal tissues were digested, RNA and DNA isolation were carried out as per Trizol manufacturer instructions.

Approximately 1-5 µg of RNA was used to create cDNA using the SuperScript III First-Strand Synthesis System for RT-PCR (Thermo 18080051). RT-qPCR was carried out using SYBR select Mastermix (Thermo 4472908) on an ABI StepOnePlus qPCR machine using the following primers: PCDH15_CD1 forward primer 5’-ctctatgaagaacttggagacagct-3’, reverse primer 5’-ggaagaaaagggcatcacaacttg-3’; PCDH15_CD2 forward primer 5’-ctctatgaagaacttggagacagct-3’, reverse primer 5’-cctcactaggctctctaatttcaactt-3’; PCDH15_CD3 forward primer 5’-ctctatgaagaacttggagacagct-3’, reverse primer 5’-ctcgatctacaactaacttgatcattct-3’. Expression of PCDH15 CD1 isoforms was measured via ΔΔCt of the most prominent isoform in comparison to GAPDH or RPS19 calibrator genes.

### AAV round window membrane injection in neonatal mice

P1 pups were anesthetized using cryoanesthesia and kept on an ice pack during the procedure. Injections were done through the RWM as previously described ^41^. A small incision was made beneath the external ear. The round window niche was identified visually, and the viral vector solution was delivered via a micropipette needle at a controlled rate of 150 nanoliters per minute. The surgical incision was closed using two sutures with a 7-0 Vycril surgical suture. After the injection, standard postoperative care protocols were implemented.

### ABR and DPOAE testing

ABRs and DPOAEs were recorded following established procedures ^61^, utilizing a custom acoustic system developed by Massachusetts Eye and Ear, Boston, MA, USA. Adult mice age 5 weeks were administered anesthesia using a ketamine/xylazine mixture and were placed on a 37 °C heating pad throughout the recording session. For ABR recordings, three subdermal needle electrodes were used: a reference electrode placed on the scalp between the ears, a recording electrode situated just behind the pinna, and a ground electrode positioned in the posterior region near the tail. Tone-pip stimuli with a duration of 5 milliseconds and a rise-fall time of 0.5 milliseconds were delivered at frequencies ranging from 4.0 kHz to 32 kHz, with alternating polarity, at a rate of 30 stimuli per second. The recorded responses were subjected to amplification (×10,000), band-pass filtering (0.3–3 kHz), and averaging (×512) using a PC-based data acquisition system running the Cochlear Function Test Suite software package provided by Massachusetts Eye and Ear, Boston, MA, USA. Sound levels were incremented in 5-dB increments, starting from approximately 20 dB sound pressure level (SPL) and increasing up to 80 dB.

The ABR Peak Analysis software (version 1.1.1.9, Massachusetts Eye and Ear, Boston, MA, USA) was used to determine ABR thresholds and measure peak amplitudes. ABR thresholds were verified visually as the lowest stimulus level at which a repeatable waveform could be discerned. DPOAEs were recorded for primary tones with a frequency ratio of f2/f1 = 1.2, where L1 was set as L2 + 10 dB. The f2 frequency ranged from 5.6 kHz to 32 kHz in half-octave increments. Primary tone levels were adjusted in 5-dB increments, spanning from 10 dB SPL to 70 dB SPL for f2. DPOAE threshold values were calculated from the average spectra and were defined as the f1 level required to generate a DPOAE of 5 dB SPL.

### Open Field Test

We used a 37-cm-diameter arena with consistent, low-level illumination for our experiments. The testing took place when the animals were age P35. Each animal was positioned at the side of the arena and position was recorded for a duration of 4 min. To prevent any potential olfactory distractions, the arena was thoroughly cleaned between test sessions. Video footage was subsequently analyzed using ImageJ software, and open field path outlines were generated. During the 4-minute observation period, we quantified the number of full-circle rotations, which encompassed both clockwise and counterclockwise turns.

### Rotarod and swimming tests

The rotarod test was conducted over two days. On the first day, mice were positioned within an enclosed housing on a rotating rod, initially spinning at a constant rate of 4 revolutions per minute (rpm) for 5 min. In the event of a mouse falling during this training session, they were promptly placed back on the rotating rod. On the second day, the trained mice were once again placed on the spinning rod, but this time with a start speed of 4 rpm and acceleration rate of 20 rpm/min. The duration each animal managed to remain on the device before descending to the machine floor of the housing was monitored by a timer and recorded after each trial. A 5-minute resting interval was enforced between trials, and a total of five trials were conducted for each mouse. The latency to fall off the rotarod was recorded.

In the swimming test, the mice were placed in a tank filled with water, which forced them to swim. The time mice could swim before needing rescue was recorded.

### FM1-43 loading in adult cochlea

Adult mice were subjected to anesthesia using isoflurane through an open drop method and were subsequently euthanized by cervical dislocation followed by decapitation. The otic capsules from the mice were carefully extracted and then placed in Leibovitz’s L-15 medium. Utilizing a stereomicroscope, the apical and mid-apical regions of the cochlea were microdissected. The tectorial membrane was gently pulled away to expose the sensory epithelium. To facilitate the experimentation, a solution of FM1-43 (2 µM in L-15) was directly applied to the exposed epithelium at room temperature and left on for 1 min. Subsequently, a solution of SCAS (0.2 mM) was applied. Imaging of the organs of Corti was carried out using an Olympus upright FV1000 confocal microscope equipped with a 60X1.1 NA water-dipping objective lens.

### Immunofluorescence labeling of mouse cochleas and HEK 293 cells

Adult mice were anesthetized using the isoflurane open-drop method and then humanely euthanized by cervical dislocation followed by decapitation. The cochleas were carefully dissected in L-15 medium and immediately fixed with 4% formaldehyde in HBSS for one hour at room temperature. Afterward, they were washed with HBSS and transferred to fresh 10% EDTA for two days to ensure complete decalcification. Once fully decalcified, the organs of Corti were microdissected and blocked with 10% donkey serum for one hour at room temperature. Subsequently, the samples were stained with an anti-HA antibody (abcam ChIP Grade, ab9110) diluted to 1:500 in 10% donkey serum and incubated overnight at room temperature, followed by multiple rinses in HBSS. The samples were then incubated in a blocking solution (10% donkey serum) for 30 minutes at room temperature. After that, they were incubated overnight at room temperature with a donkey anti-rabbit IgG secondary antibody conjugated to Alexa Fluor 594, diluted to 1:500 in the blocking solution, which also included Alexa Fluor 405 phalloidin to label actin. Following the secondary antibody steps, the samples underwent several rinses in HBSS and were mounted on Colorfrost glass slides (Thermo Fisher Scientific) using Prolong Gold Antifade mounting medium (Thermo Fisher Scientific). The slides were placed horizontally in the dark for 24 hr at room temperature to allow for drying before imaging.

The imaging was conducted using a Nikon Ti2 inverted spinning disk confocal microscope with Nikon Elements Acquisition Software AR 5.02, utilizing the following objectives: a Plan Apo λ 100×/1.45 oil, a Plan Fluor λ 40×/1.3 oil, and a Plan Apo λ 60x/1.4 oil.

For immunocytochemistry in HEK 293 cells, transfected cells underwent the following procedure: they were fixed with 4% formaldehyde in Hank’s balanced salt solution (HBSS) for 1 hr, then washed three times with HBSS, and subsequently blocked with 10% donkey serum for 2 hr at room temperature. We utilized either a rabbit polyclonal anti-PCDH15 (DC 811) antibody or rabbit anti-HA C29F4 antibody (#3724, Cell Signaling), both diluted to 1:200 in 10% donkey serum, and incubated them for 24 hr at room temperature, followed by several rinses in HBSS. Next, the samples were incubated in a blocking solution for 30 min and then incubated overnight at room temperature with a donkey anti-rabbit immunoglobulin G (IgG) secondary antibody conjugated to Alexa Fluor 594, diluted to 1:500 in the blocking solution. After the secondary antibody incubation, the samples underwent several rinses in HBSS and were mounted on Colorfrost glass slides (Thermo Fisher Scientific) using Prolong Gold Antifade mounting medium (Thermo Fisher Scientific). Imaging was carried out using an Olympus FluoView 1000 confocal microscope equipped with a 60×/1.42-NA oil-immersion objective.

### Immunofluorescence labeling of NHP retina and human retina organoids

The nonhuman primate was euthanized at 65 days after vector injection and was perfused with heparinized saline followed by 4% formaldehyde. Collected eye globes were postfixed for another 24 hr in 4% formaldehyde before retinas were dissected.

Retina samples and organoids were cryoprotected by incubating in gradient concentrations of sucrose and were embedded in OCT compound and stored at −80°C prior to sectioning. Cryosections were generated using a Leica CM 3050 S cryostat at 30-µm step size.

For immunofluorescence labeling, the following primary antibodies and secondary antibodies were used: mouse anti-HA antibody (1:100) (16B12, BioLegend), rabbit anti-HA antibody (1:200) (ab9110, Abcam), rabbit anti-ARR3 antibody (1:200) (HPA063129, Sigma Aldrich), sheep polyclonal anti-PCDH15 (1:200) (AF6729, R&D Systems), donkey anti-rabbit IgG secondary antibody conjugated to Alexa Fluor 594 (1:200), donkey anti-sheep IgG conjugated to Alexa Fluor 647 (1:200), donkey anti-mouse IgG conjugated to Alexa Fluor 488 (1:200), donkey anti-mouse IgG conjugated to Alexa Fluor 594 (1:200), and donkey anti-rabbit IgG conjugated to Alexa Fluor 488 (1:200) (Invitrogen). Samples were blocked with 10% donkey serum for 1 hr at room temperature. Antibodies were diluted in 10% donkey serum and incubated overnight at room temperature, followed by several rinses in HBSS. Next, samples were incubated in a blocking solution for 30 min and incubated overnight at room temperature with a secondary antibody in a 1:200 dilution in the blocking solution. We used DAPI or Hoechst to label cell nuclei (1:500) and BODIPY to label membranes (1:1500). Tissues were mounted on a Colorfrost glass slide (Thermo Fisher Scientific) using Prolong Gold Antifade mounting medium (Thermo Fisher Scientific). Imaging was performed with a Nikon Ti2 inverted spinning disk confocal using a Plan Fluor 40×/1.3 oil objective, and Plan Apo λ 60×/1.4 oil and a Plan Apo λ 100×/1.45 oil objective.

### Quantification of confocal microscopy data

Microscopy data analysis and quantification were done in the Fiji distribution of ImageJ v1.53. Transduction efficiency in cells was evaluated as previously described^27^. Hair cells were identified with phalloidin staining of bundles and photoreceptors with BODIPY, transduced cells by positive HA-tag labeling. Control samples without AAV were used to correct for autofluorescence. Segments with dissection-related damage were removed from the analysis.

GraphPad Prism 7 software was used to generate the graphs and perform the statistical analysis. The results are shown as mean ± SEM or mean ± SD as indicated in figure legends. Randomization was used whenever possible.

### Conventional scanning electron microscopy

Scanning electron microscopy in adult mice, NHP retina or retina organoids was performed as previously described ^62^. Immediately post-extraction, cochleas underwent pre-fixation by immersion in a solution containing 1% glutaraldehyde and 4% formaldehyde, both in 0.1 M cacodylate buffer (pH 7.2) supplemented with 2 mM CaCl_2_, for 1 hr at room temperature.

Following the pre-fixation step, the samples were postfixed using 2.5% glutaraldehyde in 0.1 M cacodylate buffer (pH 7.2), supplemented with 2 mM CaCl_2_, for an additional hour at room temperature. They were next rinsed in 0.1 M cacodylate buffer (pH 7.2) and then with distilled water. The cochlear bone was removed using a 27-gauge needle, followed by microdissection of the organ of Corti.

Next, the samples were immersed in a saturated aqueous solution containing 1% osmium tetroxide for an hour in a light-controlled environment, and then were postfix-fixed using a 1% tannic acid aqueous solution for an hour in the dark. Finally, the samples underwent a series of rinsing, dehydration, and critical point drying. The prepared samples were mounted onto aluminum stubs equipped with carbon-conductive tabs. They were then sputter-coated with a platinum layer to a thickness of 5 nm using an EMS 300 T dual-head sputter coater and imaged with an Hitachi S-4700 scanning electron microscope.

Immunogold scanning electron microscopy in mouse cochlea and human retina organoids Immunogold scanning electron microscopy was conducted in accordance with previously described protocols ^27,62^. Cochleas and human retina organoids were fixed using 4% formaldehyde. Following fixation and washing, samples were subjected to a 2-hr blocking step at room temperature using 10% normal goat serum. Next, the samples were incubated with primary antibodies for 24 hr at room temperature and were subsequently rinsed three times for 10 min each in HBSS. A an anti-HA tag antibody (Abcam, ab9110) at a 1:200 dilution in 10% donkey serum was used to label the HA tag. Following rinsing, the samples were blocked for 30 min at room temperature, using 10% normal goat serum. They were then incubated overnight at room temperature with a secondary antibody solution consisting of 12-nm Colloidal Gold Affini Pure Goat Anti-Rabbit IgG (Jackson ImmunoResearch, 111-205-144) at a 1:30 dilution in the blocking solution. Following the application of the secondary antibodies, the samples underwent three 10-min rinses in HBSS.

Finally, the samples were prepared for observation. This involved a dehydration step, followed by critical-point drying, mounting, and sputter-coating with palladium in the range of 3 to 5 nm. Samples were imaged using an Hitachi S-4700 field-emission scanning electron microscopy equipped with a backscattered electron detector.

### Statistics and reproducibility

All experiments, with the exception noted below, were reproduced three independent experiments using independent samples.

FM1-43 loading experiments in Fig. 3f and g and immunofluorescence analysis in Fig.3c and d were reproduced in four independent experiments.

Only samples displaying significant dissection-related damage were excluded from the analysis. No other data were omitted.

All figures were created using Adobe Illustrator 2022 (v26.3.1, Adobe, San Jose, CA, USA). The ABR data were organized and calculated using Microsoft Excel 2016 (version 16.0.5378.1000, Microsoft, Redmond, WA, USA) before significance calculations. Graphs were generated and statistical analysis was conducted using GraphPad Prism 7 software (version 7.04, GraphPad Software Inc., Boston, MA, USA). The results are presented as either mean ± SEM or mean ± SD, as specified in the figure legends. Randomization was employed whenever possible.

## Data availability

All data generated or analyzed during this study are included in this article and its supplementary information files. Source data are provided with this paper.

## Acknowledgements

We thank Drs. Christine Petit from Institut Pasteur and Ronna Hertzano from University of Maryland School of Medicine for providing Myo15-Cre mice. We appreciate the use of the Nikon Ti2 inverted spinning disk confocal at the Harvard Medical School MicRoN Microscopy Core and the Hitachi S-4700 scanning electron microscope at the Harvard Medical School Electron Microscopy Facility. We especially thank Bruce Derfler for laboratory management. This work was supported by a National Institutes of Health grant R01-DC016932, the Bertarelli Foundation, the Usher 1F Collaborative, the Seamans Family (to D.P.C.), the Swiss National Science Foundation (grants PCEFP3_202756 and 310030_192665, B.G.) and an NIGMS Training in Genetics Fellowship T32-GM007748 (to K.T.B.).

## Author contributions

M.V.I. conceptualization, data acquisition, data analysis, data interpretation, visualization of *in vivo* and *in vitro* experiments, manuscript writing; D.M.H. plasmid cloning, data acquisition, data analysis and data interpretation of ABR, RT-PCR experiments and some *in vitro* experiments, manuscript writing; E.M.M. plasmid cloning, data acquisition, data analysis and data interpretation of *in vitro* experiments; K.T.B. RT-PCR experiments, disease-associated variants analysis, manuscript writing; M.W. data acquisition and data analysis of human retina organoids experiments; A.J.K. data acquisition of ABR experiments; X.C. data acquisition, data analysis of NHP experiments, Y.L. neonatal injections; C.W.P. generation of R245X mouse, RT-PCR experiments in NHP; B.G. supervision, data interpretation of human retina organoids experiments, manuscript writing; D.P.C. conceptualization, supervision, data interpretation, manuscript writing.

---

**Supplementary Fig. 1.**
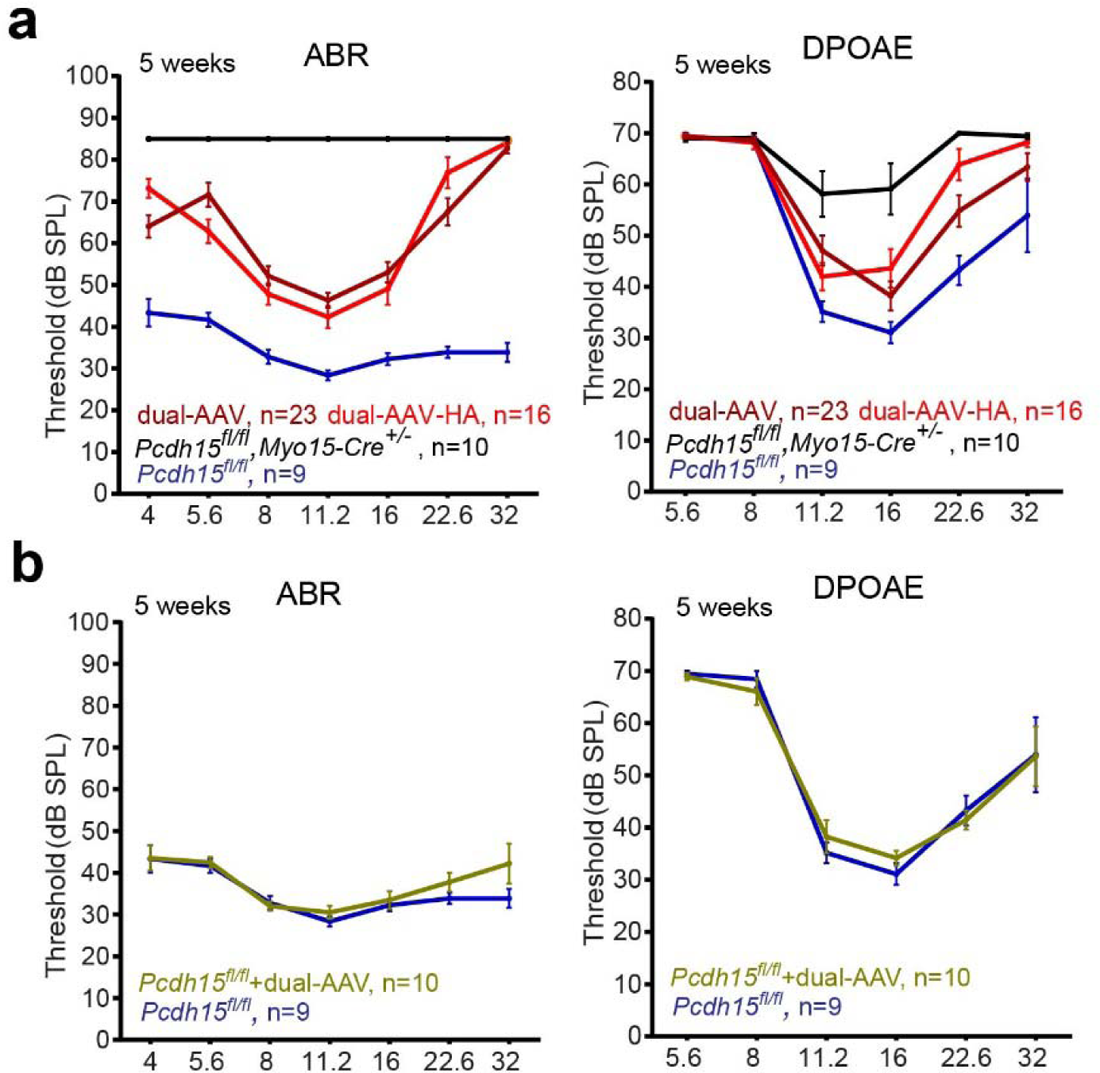
ABR and DPOAE recording in *Pcdh15^fl/fl^* Cre-hearing control mice injected with dual AAVs (n=10) demonstrated normal hearing at all frequencies, similar to uninjected *Pcdh15^fl/fl^* control mice (n=9), indicating no vector toxicity. Data are presented as mean ± SEM. Source data are provided as a Source Data file.

**Supplementary Fig. 2.**
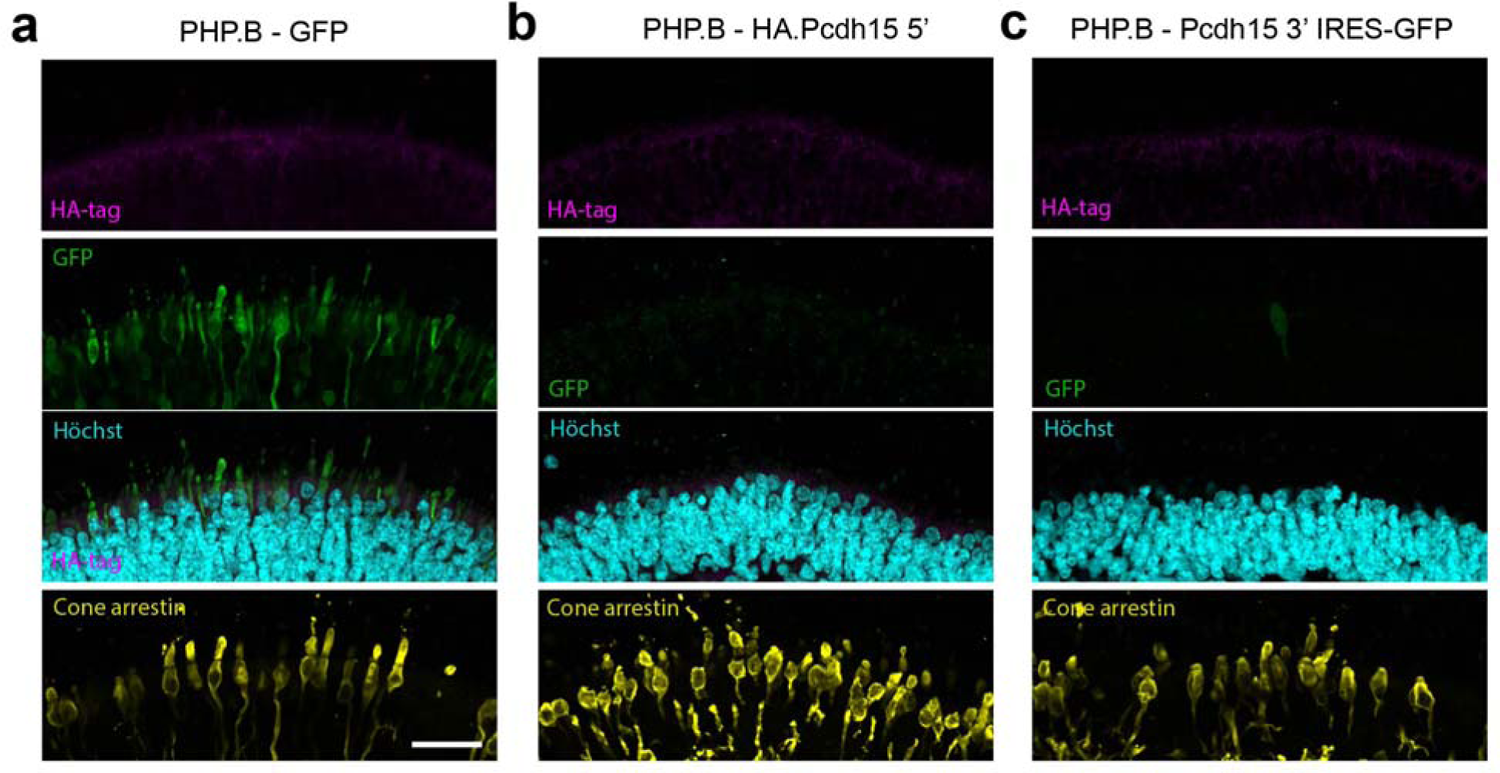
**a** Transduction efficiency of AAV9-PHP.B-CMV-GFP in human retinal organoids. **b,c** No HA tag detected when organoids were transduced with either AAV-HA.PCDH15 5’ or AAV-PCDH15 3’

